# Local control of cellular proliferation underlies neuromast regeneration in zebrafish

**DOI:** 10.1101/2024.10.22.619208

**Authors:** Natalia G. Lavalle, Jerónimo Miranda-Rodríguez, Emanuel Cura Costa, Augusto Borges, Oriol Viader-Llargués, Hernán López-Schier, Osvaldo Chara

## Abstract

Biological systems are never in equilibrium but maintain stability despite continuous external disturbances. A prime example of this is organ regeneration, where, despite intrinsically stochastic damage, organs are rebuilt through controlled cellular proliferation. In this study, we employ a cell-based computational modelling approach to investigate the proliferative response to injury. We developed a minimal two-dimensional Cellular Potts Model (CPM) using empirical data from regenerating neuromasts in larval zebrafish. Remarkably, the CPM both qualitatively and quantitatively recapitulates the regenerative response of neuromasts following laser-mediated cell ablation. Assuming that cell proliferation is locally regulated by a delayed switch, mitotic activity ceases once the type-dependent number of neighbouring cells exceeds a deterministic critical threshold. An intriguing corollary of our findings is that a local negative feedback loop among identical cells may represent a general mechanism underlying organ-level proportional homeostasis.

## 1. Introduction

One outstanding question in the regeneration field is how cells can sense damage and change from homeostasis to regeneration^1^. Regenerative cellular proliferation may be controlled systemically at the organismal level, globally at the tissue level, or locally among neighbouring cells. In regenerating organisms, including hydra^2–4^, planaria^5,6^, axolotl^7,8^ and zebrafish^9^, injuries that severely disrupt tissue integrity are followed by reactive cellular proliferation and tissue re-patterning. A remarkable example of this multilevel response is found in the neuromasts of the zebrafish lateral line.

Neuromasts are discrete circular organs that consists of around 70 cells organized into a relatively simple architecture: a central group of sensory hair cells surrounded by non-sensory supporting cells. Supporting cells are further subdivided into sustentacular cells that give rise to and physically support hair cells; and peripheral mantle cells that form an enclosing ring that delimits the organ (Fig. 1 A, A’). Recent single-cell and genetic studies have shown that there are additional subpopulations within sustentacular cells^10,11^ that are spatially coordinated to produce hair cells^12,13^.

**Figure 1.**
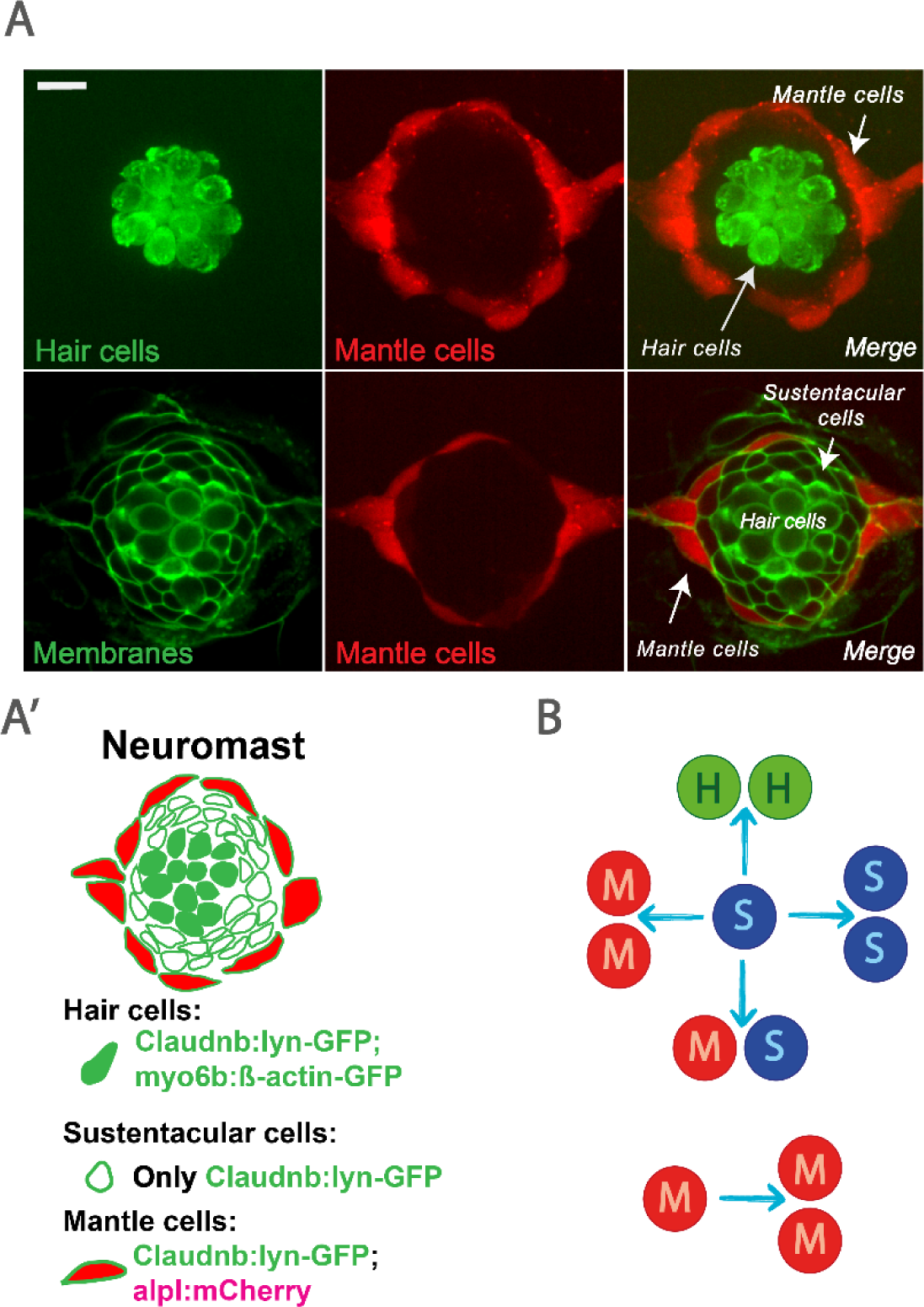
Neuromast architecture and proliferative response after laser ablation. **A)** The three main cell type classification in the neuromast organ are organized radially: hair cells are located at the center; mantle cells delimit the organ peripherally while support cells fill the space between the previous two. The cells can be distinguished in vivo by looking at fluorescent transgenic markers. The scale bar in (A) represents 10 µm. **A’)** Hair cells are uniquely marked by a β-actin-GFP fusion under transcriptional control of the myo6b promoter. Mantle cells are specifically marked by the red fluorescent protein mCherry in the alpl:mCherry transgenic. Last, the transgenic Cldb:lyn-EGFP marks the membranes of all three cell types in the organ. A triple transgenic allows to segment the membranes of all cells and to unequivocally determine the cell type in vivo. **B)** Cartoon of cell divisions in laser-ablated neuromasts (where all but 4-10 cells were ablated). 78% of sustentacular cell divisions demonstrate self-renewal, while 16% produce a pair of hair cells. In 3% of cases, they generate a pair of mantle cells, while in another 3%, one mantle cell and one sustentacular cell are generated. Cell divisions of mantle cells show complete self-renewal^9^.

Much research in the neuromast model has focused on hair cell regeneration, due to the biomedical importance of this capacity, since the hair cells of the fish lateral line are homologous to the sensory hair cells in our inner ears ^14,15^. Support cells can replenish not only lost hair cells, but also the entire neuromast when significant damage occurs. It has been experimentally determined that following the loss of up to 90% of their cells (after laser ablation of all but 4-10 cells), neuromasts regenerate by attaining radial symmetry within 3 days post-injury (dpi) and subsequently achieve a normal composition and proportions of cell types by 7 days post-injury^9^ (Fig. 1B). How different neuromast cells initiate and cease their mitotic activity remains unclear. Here, we address this question by using cell-based modelling integrating *in vivo* cellular and supracellular behaviour in regenerating neuromasts of larval zebrafish.

By developing a Cellular Potts Model (CPM), we tested the hypothesis that the neuromast-wide regenerative response is an emergent property of local cell-cell interactions. Comparing theoretical simulations with experimental data reveals that inner sustentacular cells, the main drivers of organ regeneration, cease to proliferate when they become surrounded by three or more cells of the same type, whereas mantle cells arrest divisions when confronted by between zero and two neighbouring mantle cells, indicating cooperative control of cell proliferation.

## 2. Results

### 2.1 Cell-based model of the zebrafish neuromast: a stable initial condition

Severe injury of zebrafish neuromasts that removes up to 95% of their cellular content is followed by an impressive regenerative response driven by the proliferation of the remaining mantle and sustentacular cells^9^. To qualitatively and quantitatively investigate the mechanisms responsible for this regenerative response, we conceived a two-dimensional cell-based computational model of the neuromast. We opted for the Cellular Potts Model (CPM) formalism, which is lattice-based and allows realistic cellular shapes and dynamics (see Computational Methods section 4.1). This model is optimal because the neuromast is a non-stratified epithelium, whose cells divide largely within a single plane perpendicular to their apicobasal axis. We modelled the uninjured neuromast as a two-dimensional disc of cells, respecting the empirical number of hair, sustentacular and mantle cells, as well as their spatial distribution. We parameterized the cell target areas and perimeters (see details in Computational Methods section 4.1) together with the corresponding Lagrange multipliers of the CPM Hamiltonian (Table S2) to reproduce on average the geometrical features of the neuromast cells (Fig. 2 top). Next, we examined the conditions under which the spatial organization of the in silico neuromast remains consistent over time by exploring the Hamiltonian parameters that account for the local binding energies (*J_ij_*) between cells and between a cell and its outside tissue (where *i* and *j* are either two neighbouring neuromast cells or a neuromast cell and the outside tissue). In particular, all the binding energies are *J_HH_*, *J_SS_*, *J_MM_*, *J_HS_*, *J_HM_*, *J_SM_*, *J_HO_*, *J_SO_* and *J_MO_*, corresponding to the binding between hair cells, sustentacular cells, mantle cells, hair with sustentacular cells, hair with mantle cells, sustentacular with mantle cells, hair cells with the outside tissue, sustentacular cells with the outside tissue and mantle cells with the outside tissue, respectively.

**Figure 2.**
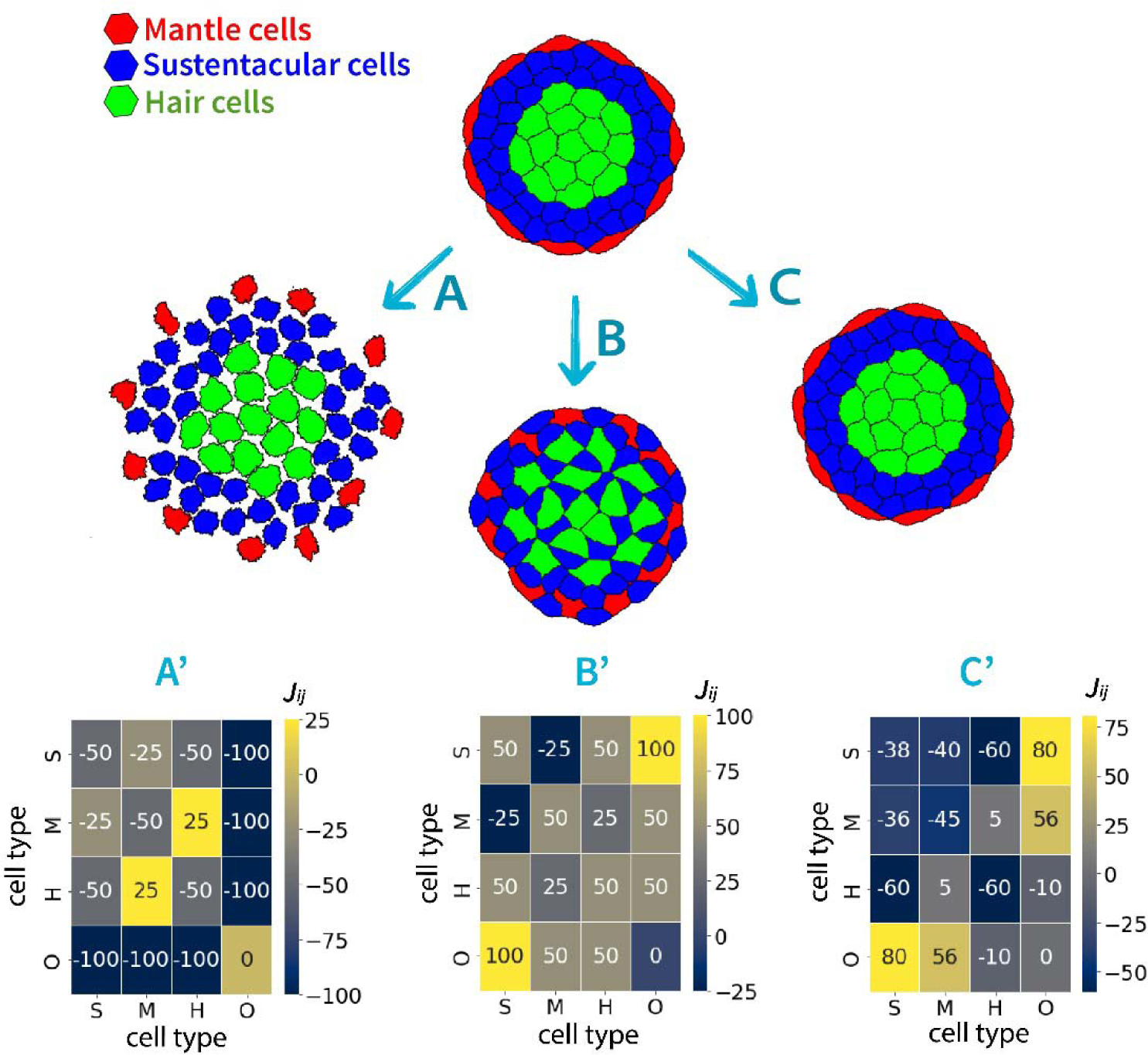
Tuning binding energy parameters leads to a stable *in silico* neuromast. (A) Non appropriate CPM binding parameter values distort the initial tissue architectures leading to disintegration or (B) anomalous tissue architectures that show intercalation of the sustentacular, hair, and mantle cells. (C) A well-defined model parameter space region can maintain stable neuromasts (Movies S1-S3). Panels A, B and C show the simulated tissues after a simulation time of 30000 MCS (Monte Carlo steps) that correspond to 1 day. Hair, sustentacular and mantle cells in green, blue and red, respectively. (A’ - C’) Heatmap displaying the corresponding energy binding parameter *J_ij_*, where the identity of the cell i and j are in the x-axis and y-axis, respectively. The specific *J* values are also listed in Table S3, while the remaining CPM parameters are detailed in Table S2.

We first excluded binding energies that, while maintaining the organ’s circular shape, strongly reduced intercellular adhesion (Fig. 2A, Movie S1 and Table S3) and increased intercalation of sustentacular with hair and mantle cells (due to a high repulsion of cells of the same type, Fig. 2B, Movie S2 and Table S3). We eventually found a convenient parametrization satisfying a stable tissue (Fig. 2C, Movie S3 and Table S3). This stable configuration was achieved by i) repulsion of hair cells with mantle cells to ensure separation (*J_HM_*> 0), ii) adhesion of sustentacular cells with both mantle (*J_SM_*< 0) and hair cells (*J_SH_* < 0), iii) repulsion of the outside tissue with the hair cells, which in turn was higher than the repulsion with the sustentacular cells (*J_OH_* > *J_OS_*> 0), and iv) adhesion of the supporting cells (sustentacular and mantle cells) higher than the adhesion of the hair cells, which in turn was higher than the homotypic adhesion of the hair cells (*J_SS_* < *J_SH_*< *J_HH_* < 0, *J_MM_* < *J_SH_*< *J_HH_* < 0, *J_SM_* < *J_SH_* < *J_HH_* < 0), in agreement with the mechanical stresses of these cells recently estimated using ForSys^16^(Table S3).

### 2.2 Uncontrolled supporting cell proliferation in the neuromast can drive unlimited neuromast growth *in silico*

Following the loss of up to 90% of their cellular content, neuromasts regenerate with a normal composition and proportions of cell types by 7 dpi^9^. Surviving mantle cells divide to produce more mantle cells. By contrast, sustentacular-cell proliferation produces offspring of all three identities. With biased probabilities, sustentacular cells divide symmetrically to produce two sustentacular, two hair cells, two mantles, or asymmetrically to give rise to one sustentacular and one mantle cell^16^. Therefore, the outcome of a sustentacular-cell division can be expressed as (Fig. 1B):

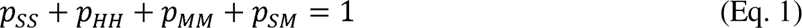

Where *p_SS_*, *p_HH_*, *p_MM_*and *p_SM_* are the probabilities that after a division, a sustentacular cell gives rise to two sustentacular cells, two hair cells, two mantle cells, or one sustentacular and one mantle cell, respectively. To recapitulate this natural response, we introduced cell divisions in the model such that the cell fates of the sustentacular cell daughters satisfying Eq. 1 (Table S1,^9^).

The position of the dividing cells correlates with the outcome of the regenerative process: sustentacular cells leading to one sustentacular and one mantle cell or two mantle cells consistently occur at the outer perimeter of the neuromast, whereas divisions generating hair cells are observed near the centre and self-renewing sustentacular cell divisions occupy the area between them^9^. This indicates that sustentacular cell fate decisions vary along the radial coordinate within the neuromast. This mediolateral correlation could reflect a signal originated at the centre of the organ or in the surrounding tissue diffusing radially out or in, respectively (see Discussion section). Thus, we modelled the dynamics of cell proliferation for the sustentacular cells by using a minimal radial potential constituted by a radial step function (Supp. Fig. S1 A). This allowed us to distinguish two regions defined by a critical distance *d_c_* with respect to the centre of mass of the neuromast. If the centre of a cell is located within a radius smaller than *d_c_,* we call them central; if it is located within a radius greater than *d_c_,* we call them peripheral. We assumed that mitosis of sustentacular cells whose centroids are within a certain critical distance *d_c_* from the neuromast centre of masses would generate two sustentacular cells with probability *p ^c^* or two hair cells with the previously reported proportion *p_HH_*. Otherwise, the two daughter cells are both sustentacular with probability *p_SS_ ^p^*. In this periphery region, cell division can also lead to two mantle cells or one mantle and one sustentacular, where the probabilities of these two last cases are the already known probabilities *p_MM_* and *p_SM_* (see section 4.1). By construction, the probability of sustentacular cell divisions occurring within a circle of radius *d_c_* (*p ^c^*) and in the periphery (*p_SS_ ^p^*) are constrained to the known *p_SS_* value above indicated in this section (Table S1).

We implemented cell division as follows. When a cell divides, a line with a random slope passing through the cellular centre of mass separates the mother cell in two, creating the future daughter cells in the next time step (Supp. Fig. S1 B i, ii and iii), the nodes of the mother cell on either side of this line are allocated to two daughter cells. Although the new-born cells have approximately half of the mother cell area, the Hamiltonian area term will drive an increase in size within a few simulation steps. We set the sustentacular cell division probability to be twice as fast as that of mantle cells^9^ (Table S2).

With this parameterized cell proliferation, we conducted simulations of a stable neuromast (parameters detailed in Tables S2 and S3) consisting of 15 hair cells, 40 Sustentacular cells, and 11 mantle cells, from which all but 4-10 cells were removed, simulating the experimental ablation. Our findings indicate that the neuromast undergoes regeneration but continues to grow unabated (see Supp. Fig. S1 C and Movie S4). Unsurprisingly, the uninjured organ also grows (Supp. Fig. S1 D and Movie S5). Hence, although the model succeeds in initiating the regenerative response, it fails in slowing it down as experimentally observed. Thus, the model requires a control mechanism to regulate the proliferative response triggered by injury.

### 2.3 Feedback switch depending on the number of neighbouring cells can control ablation- derived cell proliferation in the *in silico* neuromast

We hypothesize that cell division in the neuromast is locally regulated by neighbouring cells. Under this premise, we propose that a switching mechanism arises from the local cellular neighbourhood, modulating the rate of cell division. Specifically, when the number of neighbouring cells *N_neigh_*exceeds a certain threshold, represented by a parameter we call the ’proliferation switch’ (*S*), the cell arrests in the cell cycle and stops dividing. We implemented this mechanism and analyzed neuromast evolution after *in silico* removal of all but 4–10 cells. Initially, we modelled division based on the total number of neighbouring cells, without distinguishing their type. However, this led to neuromasts that either grew excessively or became too small, with a deficiency of hair and sustentacular cells (Supp. Figure S2 and Movies S6-S8). Therefore, we introduced a key distinction: the division rate of each mantle cell and premitotic sustentacular cell depends exclusively on the number of neighbouring cells of the same type. The results showed that the neuromast engages in a strong but transient proliferative response.

We found that the proliferation switch that best describes the experimental data occurs when sustentacular and mantle cells stop dividing after acquiring three same-type neighbours (Fig. 3A and Movie S9). However, while the model is generally capable of qualitatively regenerating the organ after injury, the steady-state tissue architecture of the regenerated neuromast contains more than three times the number of mantle cells compared to the original tissue (Fig. 3A and Movie S9).

**Figure 3.**
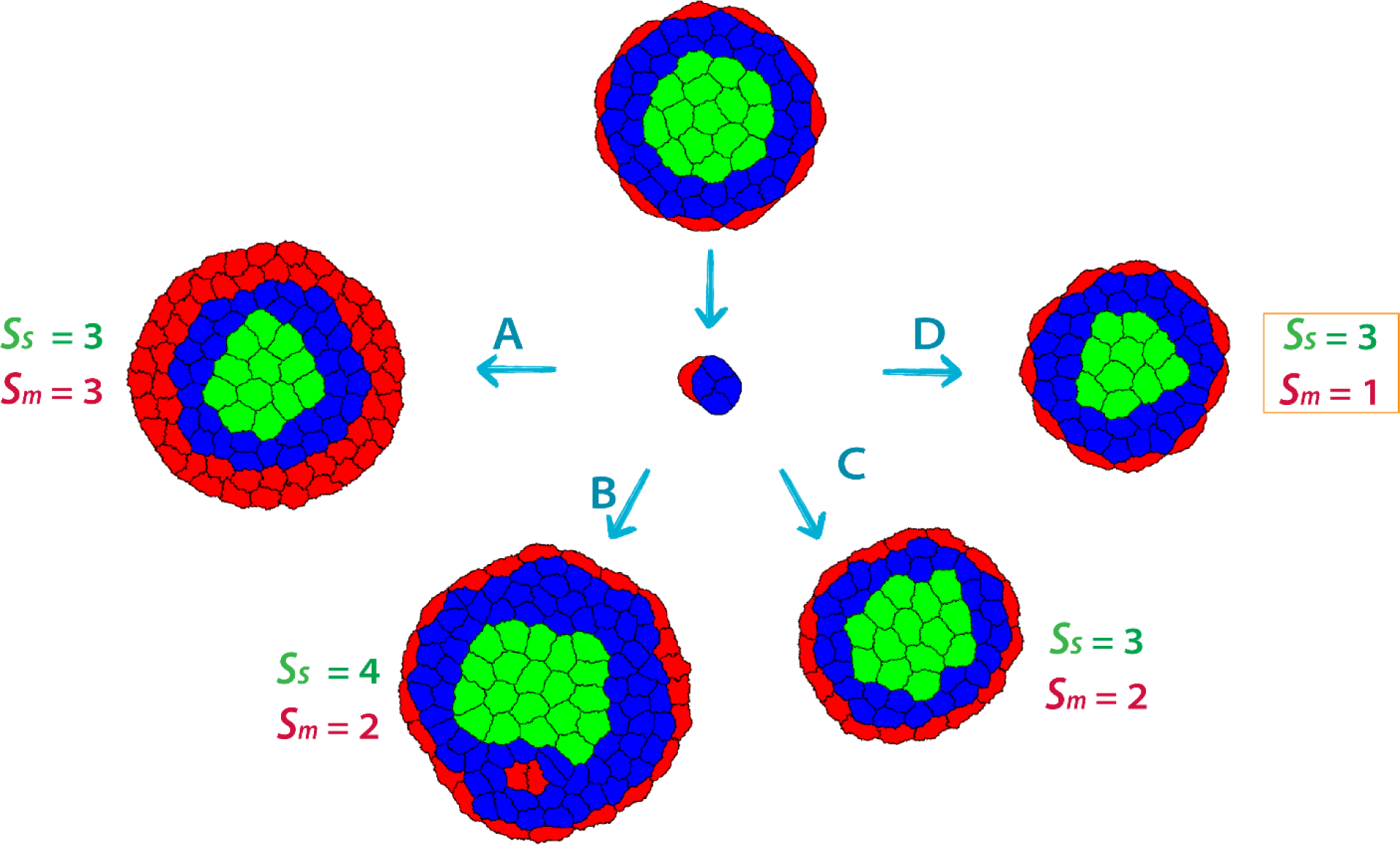
Proliferation switch of sustentacular and mantle cells control neuromast growth. (A-C) Simulation of an *in silico* neuromast subjected to an ablation in which all but 4-10 cells were removed and while the proliferation switch for mantle cells (*S_m_*) was fixed in 3(A), and 2(B and C), the proliferation of sustentacular cells (*S_s_*) was 3 (A and C) and 4 (B) .**(D)** Simulation as in (A-C) but while the proliferation switch for sustentacular cells (*S_m_*) was fixed in 3, the proliferation of mantle cells (*S_s_*) was number 1. Panel A, C and D shows the simulated tissue after a simulation time of 210000 MCS corresponding to 7 days post injury. Panel B show the simulated tissues after a simulation time of 90000 MCS corresponding to 3 days post injury. Hair, sustentacular and mantle cells in green, blue and red, respectively. All remaining model parameter values depicted in Tables S2 and S3. Note that the central tissue depicts a representative neuromast ablated leaving between 4 to 10 remaining cells. (Complete simulations depicted in Movies S7-S10)

### 2.4 Spatial heterogeneity of the feedback switch can qualitatively explain cell proliferation of supporting neuromast cells after injury

An important assumption in the previous model is that every proliferating cells have the same proliferation switch value. We hypothesized that this might explain the model’s failure to recover the original architecture of the neuromast. To address this, we developed an alternative mechanism in which sustentacular cells (s_s_) and mantle cells (s_m_) have distinct proliferation switch values, meaning that the number of neighboring cells required to halt division differs between the two cell types.

We tested this possibility by *in silico* ablating all but 4-10 cells of the neuromast and setting the cell division switch of mantle cells to s_m_ == 2. When the cell division switch of the sustentacular cells was s_s_ == 4, the ablation led to bigger and distorted neuromasts, with multiple clusters of hair cells and even transient clusters of mantle cells (Fig. 3B and Movie S10). When setting *S_s_* to three, the neuromast size and architecture begin to progress according to previously reported empirical data (Fig. 3C and Movie S11).

We next explored the cell division switch of mantle cells while fixing *S_s_* to three. Since we had already explored *S_m_*=3 and *S_m_* =2 which resulted in an excessive number of mantle cells (Fig. 3A-C and Movies S9-S11), and considering that *S_m_* =0 would imply no divisions at all, we decided to test *S_m_* =1. This parametrization qualitatively recapitulates the previously reported regenerative response of the neuromast when ablating all but 4-10 cells (Fig. 3D and Movie S12).

### 2.5 Quantitative comparison of model simulations with experimental data indicates the existence of a delay between ablation and cell division burst

To further investigate the closeness of the cell type-specific division switch to data from regenerating neuromasts, we challenged our model by comparing the predicted kinetics of total cell number and cell proportions of the simulations with the corresponding experimental data^9^. Although the results show that there is qualitative agreement between model simulations and experimental curves, we observed a distinctive difference in both, the total cell number (Fig. 4A i) and the number of each cell type (Fig. 4 A ii-iv) of the regenerating neuromast, immediately after injury. This difference suggests that cell divisions are not immediately triggered after the ablation but after a certain delay. This was confirmed empirically. Although cell death after ablation is instantaneous, cell debris remain and must be cleared by macrophages and the surviving cells must reorganize into an epithelium. Even for movies starting as early as 5 hours after injury, we observed virtually no mitosis before 20 hours after injury. During this period, we observed a conservation in the number and type of cells confirming that all cells that die do so right after ablation.

**Figure 4.**
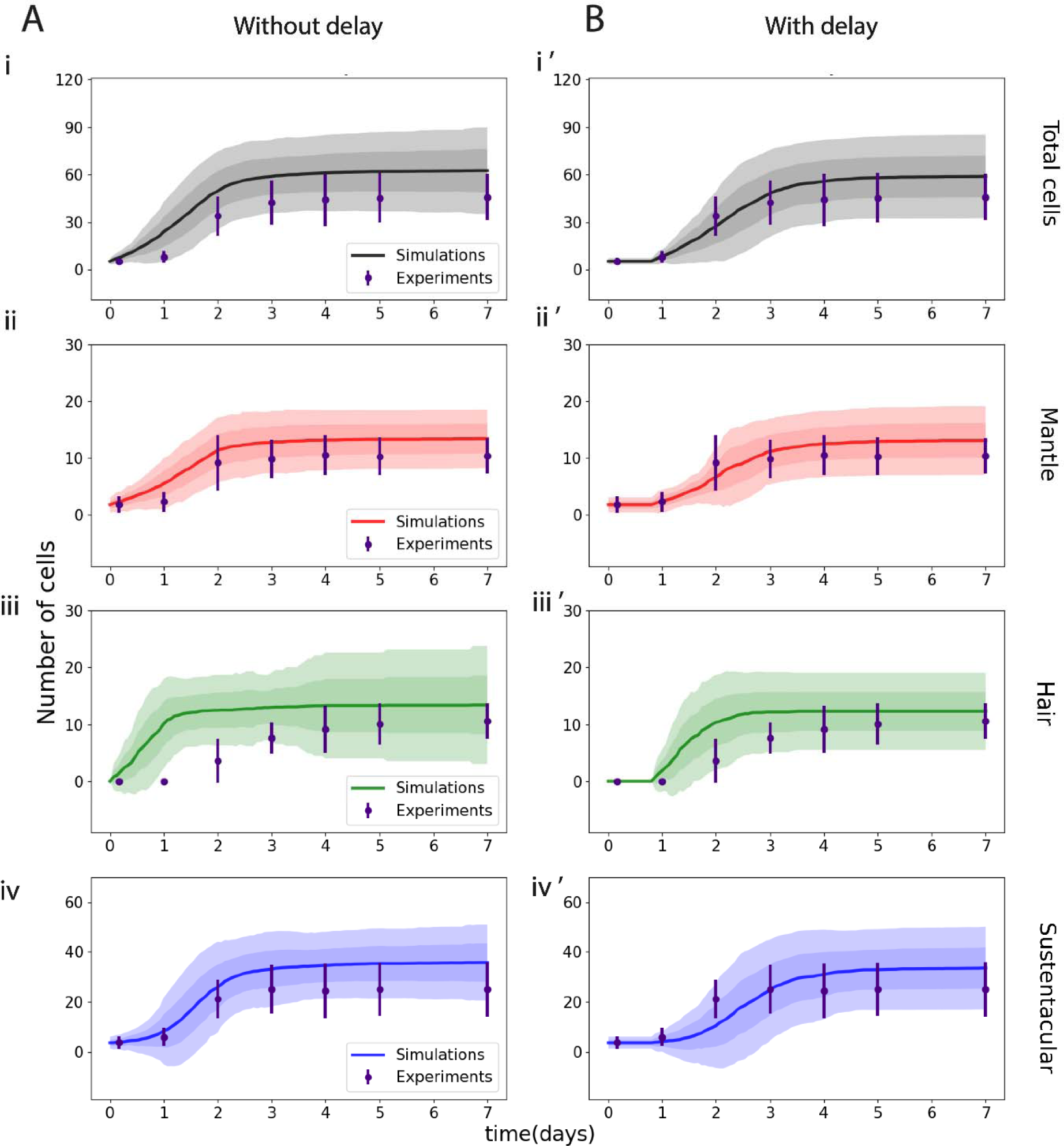
**Quantitative comparison between model-predicted simulations and experimental data suggests the existence of a delay between the ablation and cell division burst**. Predicted evolution of the neuromast cell numbers assuming that cells divide immediately after ablation **(A i, ii, iii, iv)** or after a delay of 0.8 days **(B i’, ii’, iii’, iv’)**. Time evolution of experimental and simulated neuromast total number of cells **(i, i’)** and cell proportions **(ii, ii’, iii, iii’, iv, iv’)** after an ablation of all but 4-10 cells. Cell proliferation switches are 3 for sustentacular cells and 1 for mantle cells. All model parameters depicted in Tables S2 and S3. Model simulations (with and without delay) correspond to 45 replicates obtained from 45 stochastic seeds and are depicted as shaded areas, from lighter to darker, corresponding to the 68 % and 95 % confidence intervals, respectively. Points and error bars in each panel correspond to the experimental mean and two standard deviations obtained from 15 replicates reproduced from Viader-Llargués *et a*l., 2018^9^.

To reflect this dynamic, we included an explicit delay in our model between the ablation time and the time in which cells start to proliferate, regardless of their cell type. Fitting this delay led us to conclude that the experimental data can be recapitulated when assuming a delay of 0.8 days (Fig. 4 B i’-iv’). Interestingly, this result is in agreement with the first wave of divisions previously observed^9^ (see all model parameters in Tables S2 and S3).

### 2.6 The switch on ablation-derived cell proliferation can give rise to isotropy in the neuromast regenerative response

Previous results showed that the ablation of the posterior half of the neuromast leads to a very clear regenerative response^9^. When this ablation paradigm was implemented *in silico*, our modelled neuromast was able to regenerate isotopically like its experimental counterpart. (Fig. 5 A-B, Movie S13). Furthermore, simulations in which all mantle cells were ablated resulted in a completely regenerated neuromast (Fig. 5 C and D, Movie S14), as observed in the experiments^9^. Hence, our simulations show that our model qualitatively reproduces the isotropic regenerative responses experimentally observed in the neuromast after two ablation paradigms.

**Figure 5.**
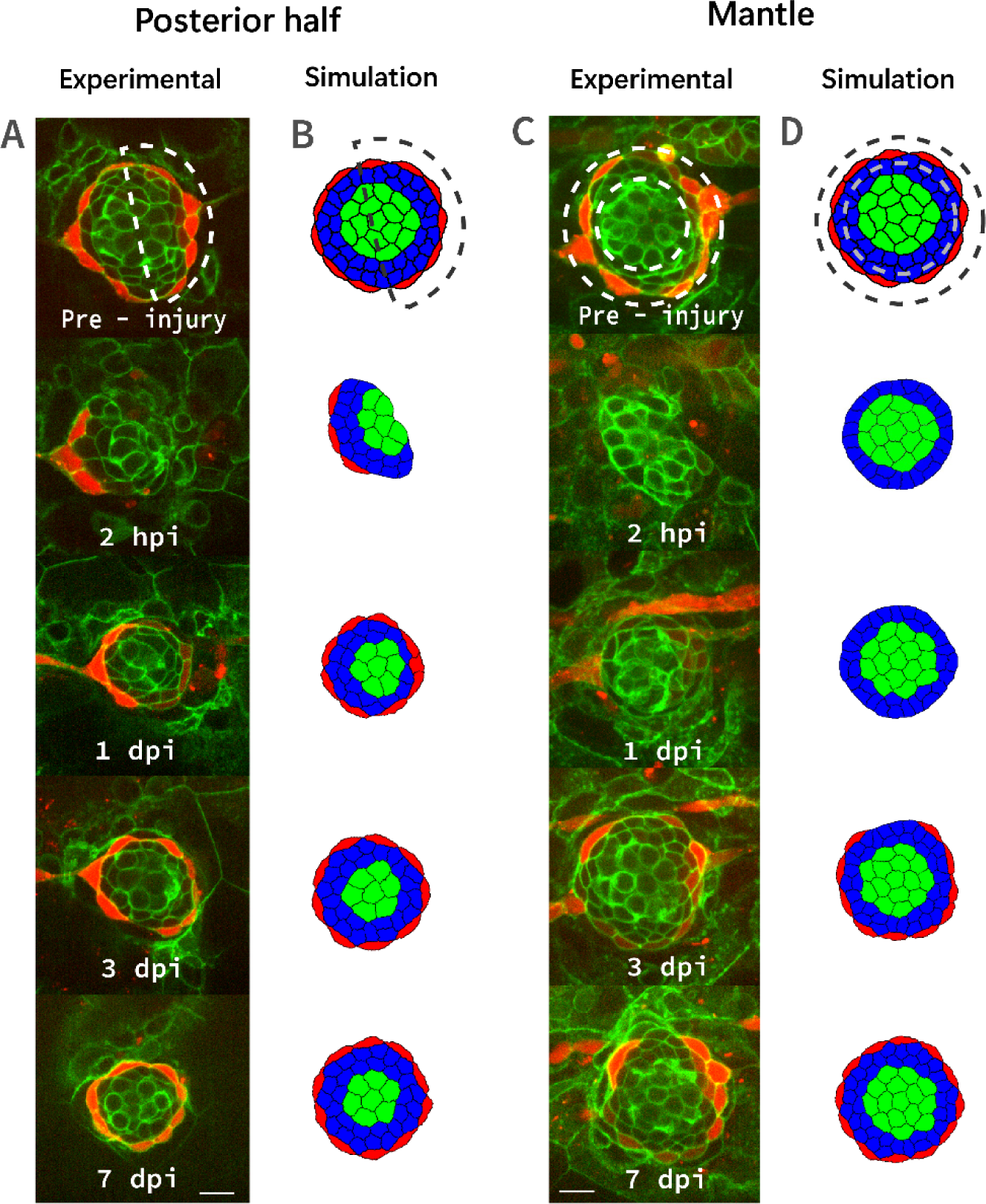
Isotropic regenerative responses of the model upon different *in silico* ablation paradigms recapitulate experimental neuromast progression during regeneration. Time evolution of experimental **(A, C)** and modelled **(B, D)** neuromasts subjected to posterior half (A, B) and mantle cells (C, D) ablation. From top to bottom, time corresponds to pre-injury, 2 hpi, 1 dpi, 3 dpi and 7 dpi in the experiments and to the initial condition, 2500, 30000, 90000 and 210000 MCS in the simulations (B, D). The two complete simulations are visualized in Movies S13 and S14. Cell proliferation switches as in Fig. 3 D. Hair, sustentacular and mantle cells in green, blue and red, respectively. CPM parameter values depicted in Tables S3 and S4. Experimental panels A and C show cell membranes in green (Cldnb:lynGFP marker) and mantle cells in red (Alpl:mCherry). The scale bar represents 10 µm.

### 2.7 The modelled proliferation switches predict neighbourhoods of supporting cells that are consistent with the experimental data

A crucial element of our model is that sustentacular and mantle cells stop proliferating when the number of neighbours of the same type reaches the corresponding switches. One would predict that sustentacular and mantle cells should have several sustentacular and mantle neighbours, respectively, similar to their proliferation switches. To test this prediction experimentally, we ablated all but 4-10 cells in the neuromasts of zebrafish larvae and detecting the number of neighbouring cells of each dividing cell. Since the regenerative response observed after ablation shows that the number of cells converges to a plateau in approximately 3 days post ablation (Fig. 4), we evaluated these predictions during this time. We ablated of cells in larval neuromasts, imaged and segmented the tissue after 3 days, and retrospectively determined cell fate after cell division to identify sustentacular and mantle cells and their neighbours 30 min before each mitosis (Fig. 6 A-E, Supp. Fig. S3, see section 4.2 for details). In parallel, we performed simulations with our model recreating the ablation of all but 4-10 cells scenario (Fig. 6 A’-D’), counted the number of neighbours of each cell type accumulated for 3 days post ablation and compared it with the experimental data. Despite the higher scatter of the experimental data, the results show that at 3 days post ablation, the number of experimental and model-predicted sustentacular cells neighbouring sustentacular cells (and the same for mantle cells) overlaps (Fig. 6 F, G).

**Figure 6.**
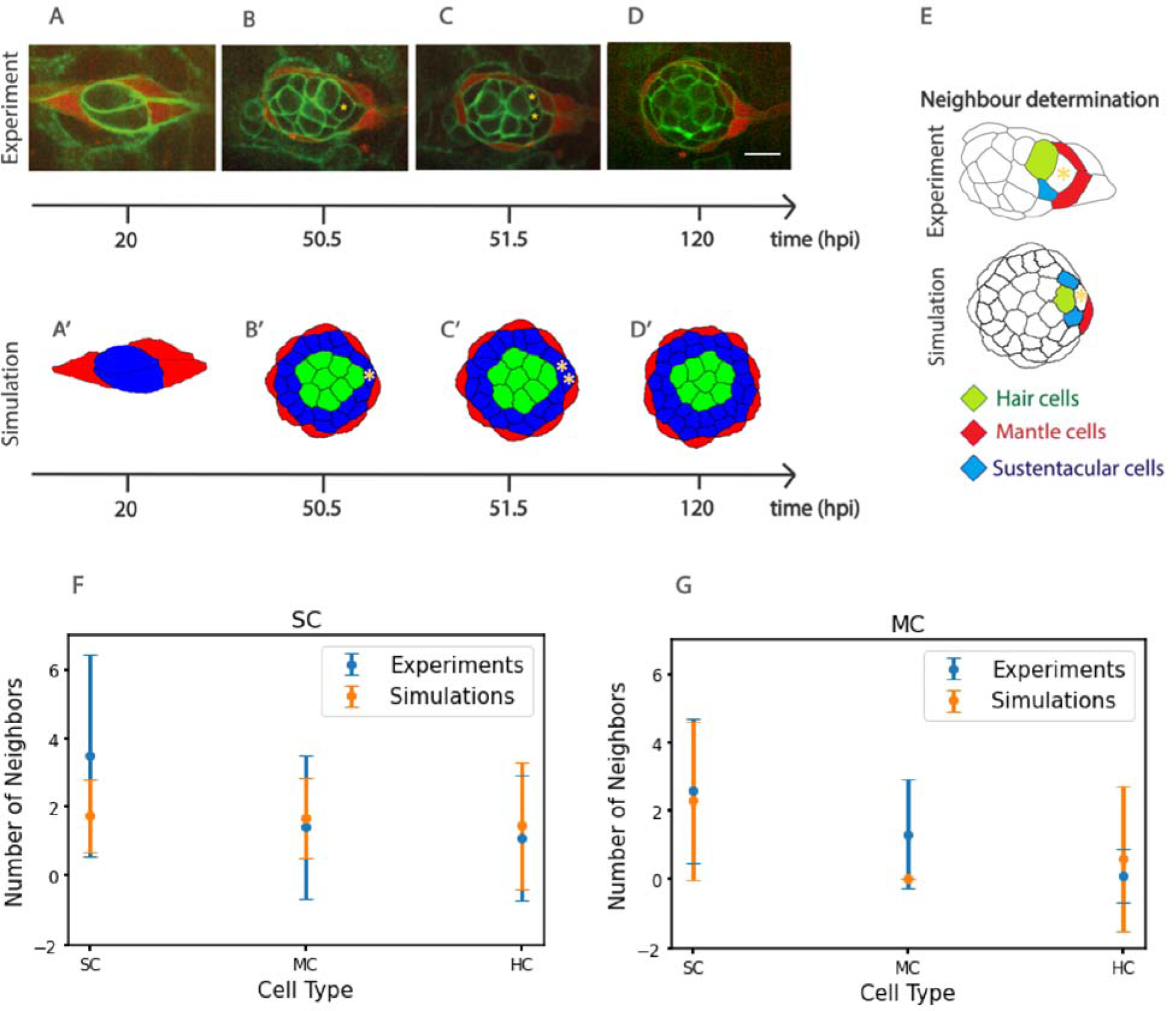
**Quantitative comparison between experimentally determined and model-predicted number of cell neighbours of neuromast cells before division**. **(A, B, C, D)** Microscopy images depicting the temporal evolution of an ablated neuromast, where the moment of sustentacular cell division is identified. **(A’, B’, C’, D’)** Simulation with our model replicating the experiment. The scale bar in D represents 10 µm **(E)** Schematic illustrating the type and number of neighbours of the sustentacular cell undergoing division (experiment above, simulation below). **(F, G)** Quantification of number of cell neighbours of sustentacular and mantle cells experimentally determined (blue) and predicted by the computational model (orange). *n* = 8 fish. The error bars correspond to two standard deviations.

## 3. Discussion

Regeneration is a tissue-scale process that channels individual cells to collectively respond to damage. In vertebrates, when the damage is severe the prevailing regenerative mechanism involves the controlled execution of cellular proliferation^17,18^. Although mitotic cell divisions are essential to replenish cell numbers, they must be tightly regulated to prevent organ hyperplasia and proportional imbalance. The neuromasts are ideally suited to construct a predictive model of regeneration because they have a well conserved size and proportions, which invariably recover during regeneration. Moreover, neuromast-size control is a purely local process that occurs independently of functional demand or systemic signals^19^.

Despite the anatomical simplicity of the neuromast, modelling regeneration of a complete organ presents various challenges. Chiefly among them is the need to integrate the cellular and supra- cellular dynamics of non-identical cells. Therefore, as a foundational approach to this problem we simulated the spatiotemporal dynamics of cells in the neuromast using a Cellular Potts Model (CPM)^20^. We found that a stable neuromast configuration emerges from a particular distribution of homo- and heterotypic adhesions and repulsions involving the hair cells and supporting cells, parameterized from previous mechanical inference performed using ForSys^16^. We could reproduce *in silico* the changes in cell numbers when introducing a delay of about 20 hours in the proliferative response to injury (Fig. 4). Remarkably, this agrees with the experimentally-observed lag between injury and the first round of divisions of the sustentacular cell population^9^.

It has been previously reported that the mitogenic activity of Wnt signalling induces sustentacular cells division in neuromasts, and that the Wnt-inhibitor Dkk limits further sustentacular-cell divisions when regenerating hair cells reach a certain threshold^21^. We reasoned that if sustentacular cells would cease proliferation by local contact with Dkk-expressing hair cells, only the sustentacular cells that are immediately adjacent to hair cells would stop proliferating, whereas more distant sustentacular cells would continue dividing. Yet, we did not see such asymmetric pattern of cell division along the mediolateral axis of the neuromast, indicating that Dkk blocks sustentacular-cell divisions equally across the tissue, or that proliferation control via Wnt occurs independently of hair cells. Therefore, in follow-up studies it would be interesting to determine sustentacular-cell numbers in neuromasts chronically lacking hair cells.

In this study, we explored an intriguing hypothesis: that the number of neighbouring cells of the same type control proliferation, which we conceptualize as akin to cell-type specific local quorum sensing^22^. By comparing model simulations with experimental data, we found that neuromast regeneration in zebrafish is consistent with sustentacular and mantle cells stopping proliferation when surrounded by three neighbouring sustentacular cells and between zero and two neighbouring mantle cells, respectively. Interestingly, this mechanism of proliferation control not only induces local maintenance of cell proportions, but also results in an emergent control of organ size.

Although the precise mechanism by which cell proliferation is halted during neuromast regeneration is beyond the scope of the present study, we speculate that it may ultimately result from mechanical cues and cell density. Interestingly, this has been observed during murine utricle development^22^. Cell fate and differential cell adhesion are related by a feedback loop, where the time of cell-cell contact promotes contact dependent signalling which in turn, provoke differential cell adhesion, because different cell types adhere to their neighbours differently^23^. During hair cell regeneration in the neuromast itself, there are examples where the cellular neighbourhood determines cell fate, such as when sustentacular cells divide to produce hair cells but only when they are located in the notch-depleted polar compartments^13^. Contact inhibition is another well-known mechanism by which cells control their proliferation rate and cell fate specification. In this regard, Notch/Emx2 signalling during hair cell formation determines the orientation of the kinocilium and the relative location of emx2+ and emx- hair cells^24,25^. Notably, our proposed mechanism of local control is consistent with the ’neighbourhood watch’ model, in which cells make decisions based only on communication with their local environment when they detect variations in signal intensity relative to their neighbours^26^. Ultimately, as spatial transcriptomics provides transcriptomic data correlating to any given cell niche, new molecular players will be discovered to participate in the cell type specific proliferation switch phenomenon^27^.

## Limitations of this study

Although our model represents the neuromast in 2D, hair cells are positioned closer to the apical surface, while sustentacular cells reside more basally^29^. This spatial organization may be relevant if the proposed local proliferation switch, based on the number of neighbouring cells, serves as an effective approximation of a more complex feedback mechanism involving the percentage of cell membrane contact between neighbours. Further studies should investigate neuromast regeneration by incorporating apicobasal organization in a 3D model to assess how this hypothetical switch is affected.

Another limitation of our model is that it does not account for the role of interneuromast cells in the regenerative response following ablation. While interneuromast cells can form neuromasts *de novo*, this primarily occurs upon loss of ErbB2 signalling^30^. Although these cells are neither necessary nor sufficient for neuromast regeneration in the laser ablation model, their absence prevents neuromasts from fully recovering their total cell number^9^. One possible explanation is that interneuromast cells regulate the target size of the organ through chemical or mechanical cues. Despite this size reduction, neuromast cell proportions and organization are restored. Thus, the proposed proliferation switch may be fine-tuned by interactions with cells outside the neuromast, particularly interneuromast cells, whose role in this process remains to be further explored.

## Conclusions

Our study demonstrates that the neuromast regenerative response is consistent with a delayed onset of cellular proliferation regulated by negative feedback from the immediate cellular neighbourhood. A key prediction of our computational model is that, on average, sustentacular and mantle cells continue dividing but stop once they have three and one neighbouring sustentacular and mantle cells, respectively. Future studies should investigate whether this mode of growth control, observed in the regeneration of the neuromast, could have broader applicability, potentially extending to other tissues and organisms and playing a role not only in regeneration but also in developmental processes.

### 4.1 Computational methods section

#### 4.1.1 Spatial Computational cell-based model of the regenerating neuromast after laser ablation

We implemented a two-dimensional Cellular Potts Model(CPM) ^31,32^ in Morpheus^33^. In this formalism, individual cells are represented as sets of sites on a discrete lattice, which in our case is a 2D grid. Each site on the lattice is identified by its position *ij* where *i* is the row and *j* the column, each cell is represented with an “ID” number (denoted as σ) and the “type” of cell it belongs to (identified by the letter τ). The ID is a unique number that identifies each cell, while the type is a label that defines the properties and characteristics of the cell, therefore, two cells have different σ but can share the same τ if they are of the same type.

In the Cellular Potts Model (CPM), when representing a system containing a total of *n* cells, each site *ij* in the lattice with the same (J, for example σ = *M* (where *M* ranges from 1 to *n*) corresponds to the cell labeled as *M*. Moreover, σ equal to 0 is utilized to denote the medium surrounding the cells.

The energy of any given cell configuration is determined by an energy function or Hamiltonian defined as follows:

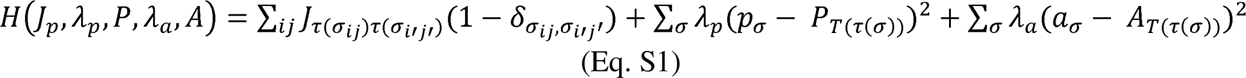

Where 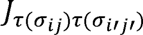 is the coupling coefficient that determines the adhesion between site ij and its selected neighbouring site *i’j*’, δ is the Kronecker delta and the second and third terms represent the energies associated with cell area elasticity and perimeter contractions, respectively, and include the Lagrange multipliers A (elasticity coefficient) and A (contractibility coefficient). “a” and “p” represent the area and perimeter of the cell σ, respectively, while “AT” and “PT” are the target area and perimeter, respectively.

The model’s evolution is based on the Metropolis algorithm3^4,35^, which involves randomly selecting a site on the lattice and evaluating whether changing the cell’s identity at that site would decrease the system’s energy. If the change results in a decrease in energy, it is accepted deterministically; otherwise, it is accepted with a probability based on the Boltzmann distribution, introducing a stochastic component. The system’s energy function (*H*) accounts for the interactions between neighbouring cells, their properties, and preferences for certain locations on the lattice. This energy function influences the stability and behaviour of cells in the tissue.

We incorporated cell division into our CPM as a probabilistic process, where at each time step, for each cell with the potential to divide (*i.e*., Sustentacular and Mantle cells), we randomly draw a number between zero and one from uniform distribution. If this number is less than a parameter, we refer to as fl for Sustentacular cells and fl for Mantle cells (Table S2), the cell undergoes division.

As described in Section 2.2, we defined a radial potential that determines two regions. To achieve this, we constructed a function ’r’ that represents the distance from the centre of a cell to the centre of the neuromast, which we defined for each cell as:

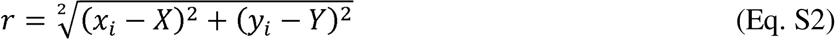

Where x_i_ corresponds to the ’x’ coordinate of the cell’s centre, X to the ’x’ coordinate of the neuromast’s centre of mass, y_i_ to the ’y’ coordinate of the cell’s centre, and Y to the ’y’ coordinate of the neuromast’s centre of mass.

Using this function, we established two regions: the central and peripheral regions, defined as r< d_C_ and r 2: d_C_, respectively, where ’d_C_’ is a scalar representing the critical distance from which we modelled cell differentiation (see Section 2.2).

The decisions regarding cell fate vary along the radial coordinate. To model the cell fate decisions of a sustentacular cell along the radial coordinate, we decided to distribute the experimentally found probabilities (Eq. 1) into these two regions (central and peripheral). We arranged it in such a way that the total divisions resulting in two hair cells occur in the central region, the total divisions resulting in mantle cells and their progeny occur in the peripheral region, and those resulting in two sustentacular cells may occur in both regions. As cell fate implementation is stochastic in our model, we constructed two probability equations, one for each region, central and periphery respectively:

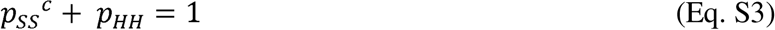

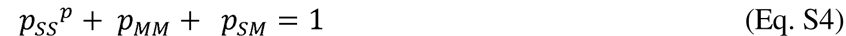

#### 4.1.2 Spatial cell-based model implementation and parametrization

We implemented our computational model using Morpheus, version 2.2.6^33^. To estimate the target areas (*A_T_*) and target perimeters (*P_T_*) for each cell type, we analyzed five epifluorescence microscopy images taken at the final time point of independent neuromast ablation experiments. Using Fiji^36^ we measured the area and perimeter of each cell in every image. Finally, we averaged the area and perimeter of all cells of the same type across all images to obtain our (*A_T_* and *P_T_*) values (Table S2).

To parameterize time, we conducted various simulations by increasing the number of Monte Carlo Steps (MCS) equivalent to one day until we found the value at which the proper tissue convergence is ensured. We found that the optimal simulation time corresponds to 30,000 MCS, equivalent to one day.

We explored different values of d_C_ and we found that values less than 80 and greater than 100 generate hypotrophy and hypertrophy in the number of hair cells, respectively. Therefore, in agreement with what was experimentally observed^9^(refer to Figure 8F in that reference), we decide to analyse five different values of d_C_ between 80 to 100: 80, 85, 90, 95, and 100 nodes, equivalent to 8.16, 8.67, 9.18, 9.69, and 10.2 micrometre, respectively (0.102 micrometres = 1 lattice nodes). For each one, we performed 30 simulations starting with 4-10 cells and compared the average final number of cells of each type with the average values from 15 experiments obtained from Viader-Llaguer et al^9^. The optimal value of d that we obtained was 95 nodes (9.69 micrometres), which is approximately equivalent to one and a half hair cell diameters, given that hair cells in zebrafish neuromasts typically range from 5 to 7 micrometres in diameter.

For the fl and fl parameters of our model, we decided to maintain the relationship reported in Viader-Llargués et al., 2018^9^, where fl = 3/2 fl (Table S2).

To parameterize the type of progeny of a Sustentacular cell according to the region in which it is located, we relate the probabilities of a division being of type SS, SH, SM, or MM (Eq.1) to those defined in equations S12 and S13, such that a percentage of the total SS divisions measured experimentally occur in the peripheral region and the rest in the central region. We studied two possible scenarios: that 60% of SS divisions occur in the peripheral region and 40% in the central region, and that 70% occur in the peripheral region and 30% in the central region. We conducted 45 simulations for each proposed scenario. Cases where the probability of SS in the central region were less than 30% and greater than 40% were discarded as they did not reproduce the experimentally observed phenomenology. This is because our model considers neighbours of the same class as a stopping mechanism, and if SS divisions occur with a high percentage, the mechanism is activated, impeding the regeneration process of the neuromast. Conversely, if the percentage is too low, in the initial divisions, all hair cells will be generated, leading to neuromast atrophy. To determine the appropriate parameterization, we compared the average number of cells of each class in the experimental final state with the simulated values. We found that the values that best reproduced the experiments involved assigning 70% of SS divisions to the peripheral region and 30% to the central region.

78% of the total divisions of a sustentacular cell during the regeneration process correspond to SS (see section 2.1 and Figure 1B). Consequently, 70% of these (54.6%) were assigned to the peripheral region, while the remaining 23.4% were designated to the central region. The rest of the percentages stay equal (HH 16%, SM 3%, MM 3%) and are allocated to their respective regions (see 4.1). In this manner, we can derive the probabilities p_SS_^C^, p_HH_, p_SS_^p^, p_MM_, p_SM_.

For the central region we have 23.4 (%SS) + 16 (%HH) = 39.4. This calculation results in p_SS_^C^ == 0.594 and p_HH_== 0.406 . In the same way, we calculated the probabilities in the peripheral region: p_SS_^p^== 0.901, p_MM_== 0.0495 and p_SM_== 0.0495.

The simulations corresponding to the 8 experiments, in which the number of neighbouring cells of neuromast cells before cell division is counted (Section 2.7), were generated with the initial condition being reconstructed from the first microscopy image taken after laser ablation in each experiment. Since the tissue is inflamed after ablation, the size of the cells in this condition is slightly bigger than the target area. To ensure that the temporal evolution from the initial condition to the target area is not abrupt, we decided to define the Lagrange multiplier associated with the area as a sigmoidal function of time (Table S2).

Using the cell count data over time and a Python script developed for this purpose, we identified instances of cell division and recorded the number of neighbours of the same class that the cell had at the moment just before division.

### 4.2 Experimental methods

#### 4.2.1 Fish maintenance and breeding

Adult zebrafish were kept in standard conditions: 27.5°C and a light/dark cycle of 14 and 10 hours. Double transgenic fish Tg(-8.0cldnb:LY-EGFP); Tg(myo6b:actb1-EGFP) were crossed with Tg(alpl:mCherry) fish. The progeny can be screened as soon as 24hpf by looking at green fluorescence in the heart and the epidermis. 50% of these double transgenic fish are alpl:mCherry positive too, and are screened by looking for red fluorescence at the retina.

#### 4.2.2. UV Laser cell ablations

Cell ablations were performed as previously described^9,37^ 3 dpf triple transgenic larvae were anesthetized and mounted laterally on their left side in 0.8% low-melting point agarose. Either the L2 or L3 neuromast was ablated by targeting all but 5-10 cells. The fish were released from the agarose and left to rest for at least 2 hours after treatment in fresh E3 medium without anaesthetic.

#### 4.2.3 Long term videomicroscopy imaging

After the rest period, the most active larvae were put in a 1:40 diluted tricaine solution and screened for nicks in the fins or tail that could signal some damage during previous manipulations. Selected fish were mounted in 0.8% agarose containing 1:40 tricaine stock solution, and covered in the same amount of anaesthetic in E3 medium after agarose solidified. Fish were imaged with a 63X water immersion objective with Zeiss Immersol W oil for up to three days. Frames were acquired serially in the green and red channels every 15 minutes. Each frame consists of a z-stack centred on the cell bodies with 6 slices above and 6 slices below, each slice separated by 1µm. E3 medium was replaced every 12 hours with fresh medium with anaesthetic.

#### 4.2.4 Image analysis

The time lapse videos were screened manually for dividing cells. For every frame where a mitotic cell was identified, the frame -2 (30 minutes before) was selected for segmentation with EpySeg^38^. This segmentation was used to count and identify the cells in contact with the cell about to divide (Supp. Fig. S3). The cell type of each neighbour was visually annotated based on morphology; the presence or absence of red fluorescence; and the presence or absence of a hair cell cuticle (marked by myo6b:act 1b-EGFP) both in the segmented frame and throughout the movie.

## Data Availability Statement

The Morpheus-based model implementations presented in Results section 2.1-2.7, the Python-based corresponding analysis and experimental data is available in zenodo: https://zenodo.org/records/13922477 ^39^.

## Acknowledgements

The authors thank Prof. Dr. Tomás S. Grigera and all the members of the Chara laboratory for valuable comments and suggestions. N.G.L was funded by Doctoral fellowships from Agencia Nacional de Promoción Científica y Tecnológica of Argentina and CONICET, Argentina. J.R.M.-R. was funded by the European Union’s Horizon 2020 research and innovation program under the HORIZON EUROPE Marie Sklodowska-Curie Actions grant agreement 840834. E.C.C was funded by a Postdoctoral fellowship from Agencia Nacional de Promoción Científica y Tecnológica of Argentina. A.B. was funded by the BMBF 01GQ1904 grant. HL-S received funding from the New York University Abu Dhabi. O.C. was funded by grants PICT-2017-2307 and PICT-2019-03828 from the Agencia Nacional de Promoción Científica y Tecnológica of Argentina and by a Biotechnology and Biological Sciences Research Council grant (grant number BB/X014908/1).

## Author contributions

N.G.L and E.C.C. implemented and simulated the Cellular Potts Models and NGL analysed the *in silico* data. J.R.M.-R., A.B. and O.V.-L. acquired the zebrafish microscopy images and analysed the corresponding results. H.L.S. supervised the experiments and edited the manuscript. O.C. conceived the project, analysed the results, secured funding, supervised N.G.L, E.C.C., and A.B., and, wrote the manuscript with contributions of all the authors.

## Supplementary Tables

**Table S1.** Probabilities of self-renewal and differentiation of sustentacular cells.

| Probability | Value |
| --- | --- |
| $p_{MM}$ | 0.03 |
| $p_{SS}$ | 0.78 |
| $p_{SM}$ | 0.03 |
| $p_{HH}$ | 0.16 |

**Table S2.** Parameters of the CPM model for all types of cells (section 4.1). A_p_and A_a_ are the multipliers of Lagrange corresponding to perimeter and area, respectively, with P_r_ and A_r_ representing the target perimeter and area, both estimates with the experimental data from this work (see section 4.2). In brackets are the types of cells. P_s_ and P_M_ are the probabilities of cell division occurring for sustentacular (mantle) cells estimated from Viader-Llargués et al., 2018^9^.

| Parameter | Value | Estimation |
| --- | --- | --- |
| *Initial number of Mantle* | 11 (*cells*) | Estimated from Viader-Llargués et al., 2018 (9) |
| *Initial number of Sustentacular* | 39 (*cells*) | Estimated from Viader-Llargués et al., 2018 (9) |
| *Initial number of Hair* | 15 (*cells*) | Estimated from Viader-Llargués et al., 2018<sup>9</sup> |
| $\lambda_a$ (*Mantle*) | $\frac{time^2}{(1000^2 + time^2)}$ (*adimensional*) | parametrization |
| $A_T$ (*Mantle*) | 1643 (*nodes*<sup>2</sup>) | Estimated with the experimental data from this work |
| $\lambda_p$ (*Mantle*) | 1 (*adimensional*) | parametrization |
| $P_T$ (*Mantle*) | 213 (*nodes*) | Estimated with the experimental data from this work |
| $\lambda_a$ (*Sustentacular*) | $\frac{time^2}{(1000^2 + time^2)}$ (*adimensional*) | parametrization |
| $A_T$ (*Sustentacular*) | 1579 (*nodes*<sup>2</sup>) | Estimated with the experimental data from this work |
| $\lambda_p$ (*Sustentacular*) | 1 (*adimensional*) | parametrization |
| $P_T$ (*Sustentacular*) | 177 (*nodes*) | Estimated with the experimental data from this work |
| $\lambda_a$ (*Hair*) | $\frac{time^2}{(1000^2 + time^2)}$ (*adimensional*) | parametrization |
| $A_T$ (*Hair*) | 2597 (*nodes*<sup>2</sup>) | Estimated with the experimental data from this work |
| $\lambda_p$ (*Hair*) | 1 (*adimensional*) | parametrization |
| $P_T$ (*Hair*) | 206 (*nodes*) | Estimated with the experimental data from this work |
| $\Omega_S$ | 0.00015 (*adimensional*) | Estimated from Viader-Llargués et al., 2018<sup>9</sup> |
| $\Omega_M$ | 0.0001 (*adimensional*) | Estimated from Viader-Llargués et al., 2018<sup>9</sup> |
| T(temperature) | 10 | parametrization |

**Table S3.** CPM interaction parameters *J* representing the local binding energies among cells or between cells and their surrounding medium. Values corresponding to. **Figs. 2 A, B and C, where the latter corresponds to the stable configuration used in all the remaining simulations of this study.**

| Interaction parameters | Figure 2A (A.U.) | Figure 2B (A.U.) | Figure 2C (A.U.) |
| --- | --- | --- | --- |
| $J_{SS}$ | -50 | 50 | -38 |
| $J_{MM}$ | -50 | 50 | -45 |
| $J_{HH}$ | -50 | 50 | -60 |
| $J_{HM}$ | 25 | 25 | 5 |
| $J_{SH}$ | -25 | -25 | -40 |
| $J_{SM}$ | -25 | -25 | -36 |
| $J_{HO}$ | -100 | 100 | 80 |
| $J_{SO}$ | -100 | 50 | 56 |
| $J_{MO}$ | -100 | 0 | -10 |

## Supplementary Figures

**Supplementary Figure S1.**
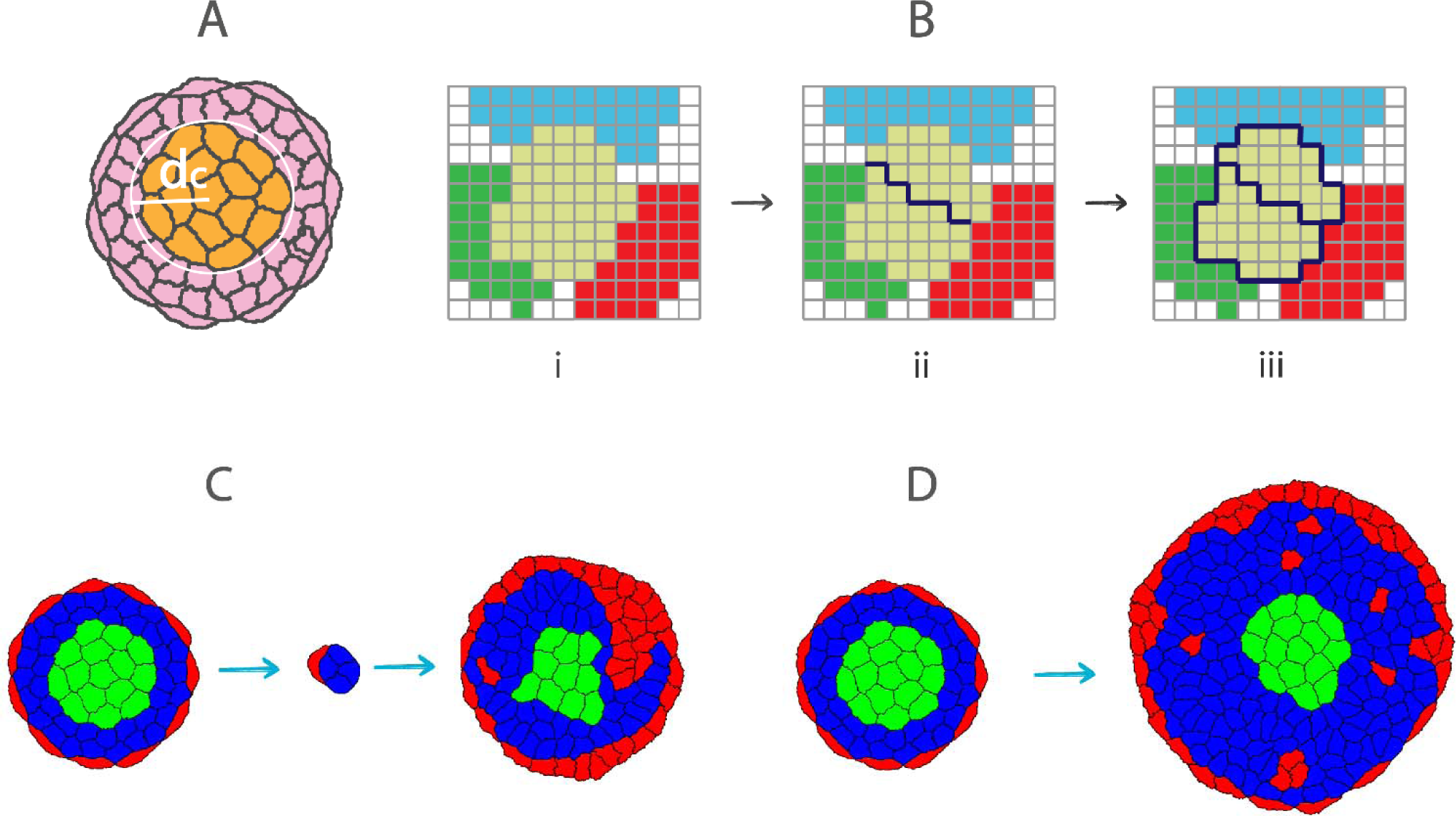
Dynamics of cell proliferation during neuromast regeneration. **A)** Schematic representation of the radial potential governing the cellular differentiation dynamics of sustentacular cells based on a critical distance to the centre of mass referred to as *d_c_* during neuromast regeneration. Within *d_c_* the sustentacular cells differentiate into two hair or two sustentacular cells. However, outside of *d_c_,* the differentiation occurs to two sustentacular, two mantle or one mantle and one sustentacular cell. **B)** Illustration of the implemented cell division process, where a line with a random slope passing through the cellular centre of mass separates the two daughter cells in the next time step. **C)** Simulation results illustrating uninterrupted growth in neuromast regeneration, initiated by retaining only 4-10 cells and simulating during 30000 MCS. **D)** in the absence of ablation, the neuromast exhibits continuous and uninterrupted growth (simulating during 90000 MCS). Parameter values of simulations depicted in C-D in Tables S2 and S3.

**Supplementary Figure S2.**
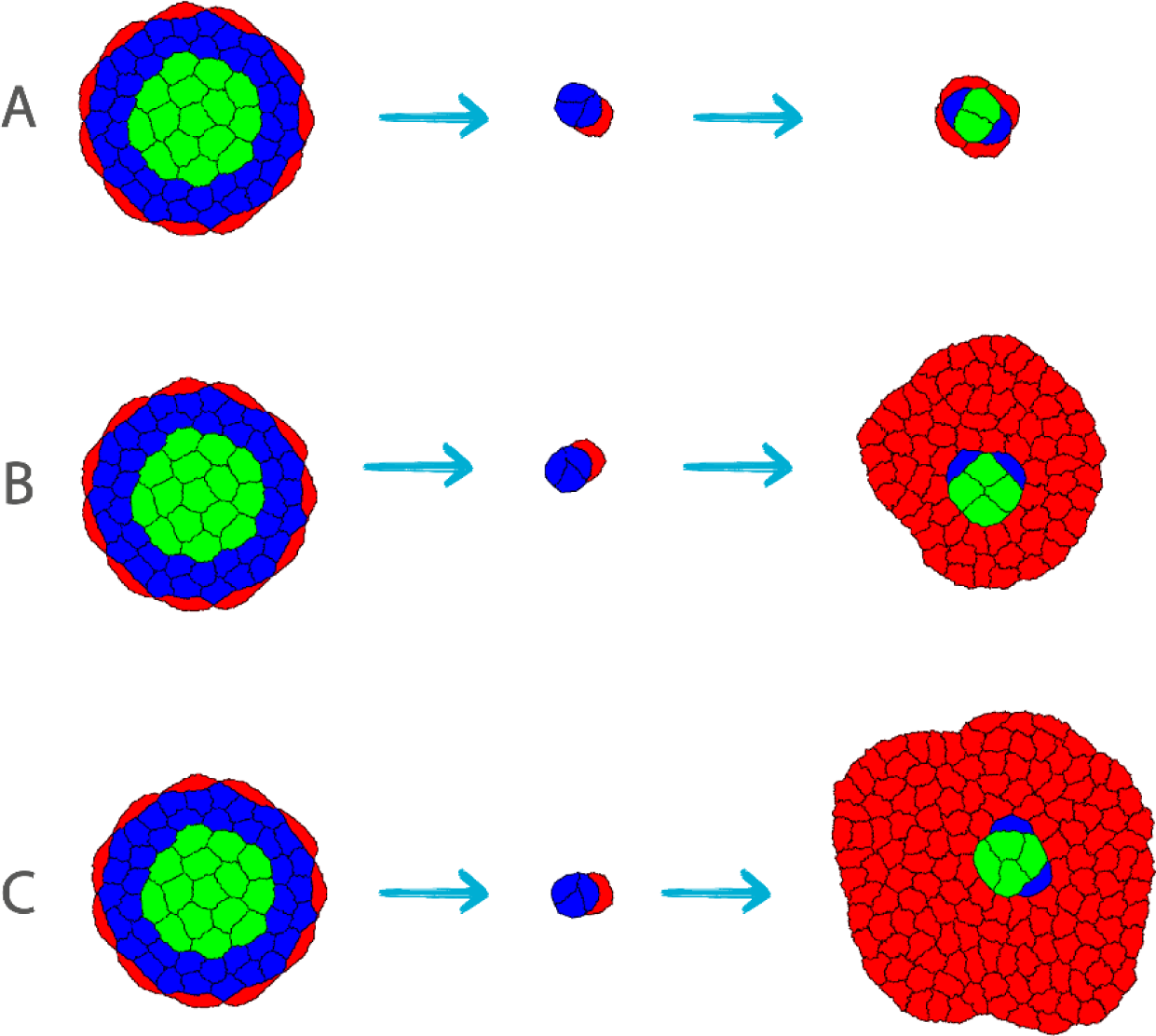
Modulation of cell division rate based on the local cellular neighbourhood, regulated by a proliferation “switch” (*S*) that depends solely on the number of neighbouring cells, regardless of their type. (A) Simulation of neuromast regeneration initiated with 4–10 cells after ablation and run for 150,000 MCS, with *S* = 3. This condition results in an atrophied neuromast with very few cells. (B) With *S* = 4, the simulation (150,000 MCS) leads to excessive growth, with an overaccumulation of mantle cells. (C) A simulation with *S* = 5, run for 80,000 MCS, results in continuous growth and an excess of mantle cells. Parameter values for the simulations shown in A–C are detailed in Tables S2 and S3.

**Supplementary Figure S3.**
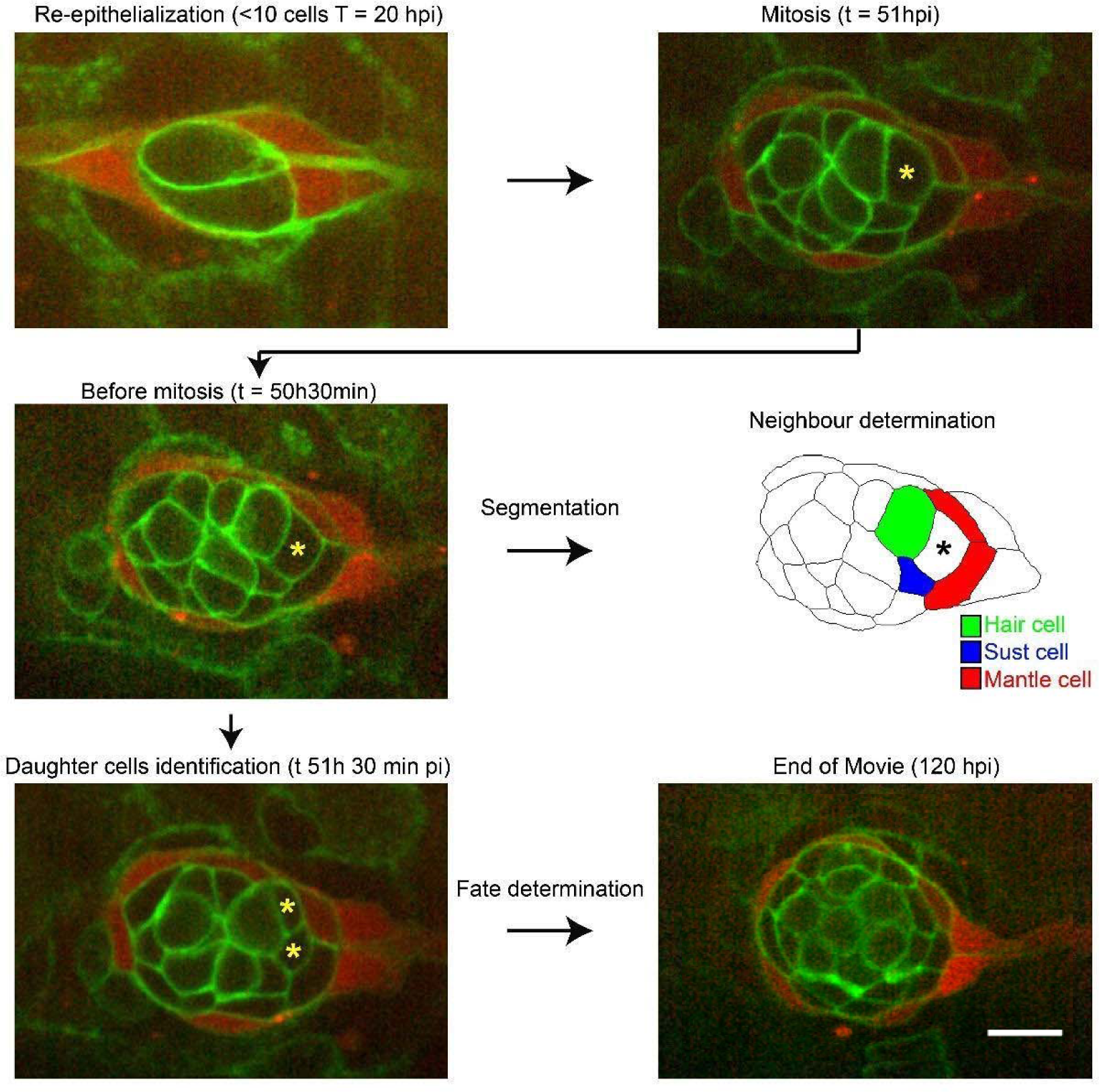
Workflow for cell neighbourhood annotation. Cell divisions do not occur in the ∼20 hpi as the debris of the death cells are cleared and the tissue re-organizes. For each mitosis in the movie, it is registered the time of division and the final cell-type of the two sibling cells based on morphology and fluorescent markers. Therefore, a dividing cell is classified as HH, SS, MS or MM based on the fate of the progeny (hair cells always appear in pairs, so no dividing cell is HS). For each division, the movie is rewound by 30 minutes, at which point, the neighbours of the dividing cell are identified, as well as their cell type and counted. The scale bar represents 10 µm.

## Legends of Supplementary Movies

**Movie S1. Model of an unstable neuromast.** Details of model parameterization described in the legend of Fig. 2 A.

**Movie S2. Model of a stable but unrealistic neuromast.** Details of the movie described in the legend of Fig. 2 B.

**Movie S3. Model of a stable and realistic neuromast.** Details of model parameterization described in the legend of Fig. 2 C.

**Movie S4. Model of a growing and regenerating neuromast.** Details of model parameterization described in the legend of Supp. Fig. 1 C.

**Movie S5. Model of a growing and uninjured neuromast.** Details of model parameterization described in the legend of Supp. Fig. 1 D.

**Movie S6. Model of a regenerating neuromast.** Details of model parameterization described in the legend of Supp. Fig. 2 A.

**Movie S7. Model of a regenerating neuromast.** Details of model parameterization described in the legend of Supp. Fig. 2 B.

**Movie S8. Model of a regenerating neuromast.** Details of model parameterization described in the legend of Supp. Fig. 2 C.

**Movie S9. Model of a regenerating neuromast.** Details of model parameterization described in the legend of Fig. 3 A.

**Movie S10. Model of a regenerating neuromast.** Details of model parameterization described in the legend of Fig. 3 B.

**Movie S11. Model of a regenerating neuromast.** Details of model parameterization described in the legend of Fig. 3 C.

**Movie S12. Model of a regenerating neuromast.** Details of model parameterization described in the legend of Fig. 3 D.

**Movie S13. Model of a regenerating neuromast after ablation of the posterior half of the neuromast.** Details of model parameterization described in the legend of Fig. 5 A-B.

**Movie S14. Model of a regenerating neuromast after mantle ablation.** Details of model parameterization described in the legend of Fig. 5 C.

